# An in-vitro investigation of effects of acetylcholine on attenuate and low motile diluted rooster semen quality

**DOI:** 10.1101/2022.05.03.490548

**Authors:** Osama. R. Ghaffar, Amjad Farzinpour, Asaad Vaziry, Sina Naderi

## Abstract

This study was conducted to evaluate the effects of acetylcholine (ACh) on 24 hours preserved diluted rooster semen at 4°C with a low motility percentage during 2 hours at room temperature incubation. Ten indigenous same-aged roosters were used in this study, adapted by Dorso-abdominal massage for semen collection for one month before starting main work. Semen was collected weekly twice by the Dorso-abdominal massage technique, collected semen transferred to the laboratory in less than 10 minutes, and took a previous qualifying examination. Qualify semen from each rooster polled together and diluted by the Lake extender and preserved for 24 hours at 4°C in the refrigerator. Different ACh concentrations (10mM-100mM) were added to Low motile semen and stored for 24 h at 4°C, and the quality parameters such as motility, viability, acrosome, and plasma membrane integrity were measured evaluated. Adding ACh in 10mM for the low motile semen significantly increased semen motility from 50% to 78.5% (P<0.05) at time zero, but other ACh concentrations did not have any significant differences compared to control. During two hours’ incubation of recovered semen at room temperature, 10mM ACh prevented declining semen motility compared to control and other ACh concentrations significantly (P<0.05), which motility in 10mM ACh concentration 59%. In contrast, control, 1mM ACh, and 100μM ACh group was 2.5%, 4%, and 1%, respectively. Semen viability after two hours of recovery at room temperature significantly 3.5% was less than the control group (P<0.05), but acrosome and plasma membrane integrity has not had any differences between all experimental groups (P>0.05). We can conclude that 10mM ACh can recover semen motility and not have toxicity and side effects on semen quality.

## Introduction

Cholinergic systems are one of the vital systems in the body’s nervous system. Neurotransmitter acetylcholine (ACh), synthesis of molecules, receptors, transporters, and enzymes for its degradation are parts of this system. Otto Loewi and Henry Dale, for the first time in 1926, discovered the ACh and, for this foundation, received a Nobel prize in 1936. After discovering ACh in the nervous system: Identified ACh as a post-ganglionic neurotransmitter in parasympathetic neurons and a preganglionic-neurotransmitter in sympathetic neurons in several parts of the brain and body [1]. According to phylogenesis, in non-neuronal cells such as monera, protozoa, algae, tubellariae, and primitive plants, fortunately, are shown the cholinergic system; for this reason, this kind of cholinergic system is called the non-neuronal cholinergic system (NNCS) [2]. Founded the NNCS in animal spermatozoa and started researching the effect and function of each compartment of this system on spermatozoa in the six-teens and seventeens decade of last century.

ACh is synthesized within the cellular cytoplasm from choline and acetyl coenzyme A (acetyl Co-A) by a molecular protein that is known as a (choline acetyltransferase enzyme (ChAT)) in neuronal and most of the non-neuronal cells [3]. Flagella are responsible for spermatozoa movements and contractile proteins [4]. Contractile proteins of flagella are generally similar to the smooth muscle, and scientists believe that the ACh cycle is responsible for the contraction and relaxation of flagella [5]. In the mammalian spermatozoa, all compartments of the cholinergic system include ACh, ChAT enzyme [6], AChE [7], and both nAChRs and mAChRs are present [5]. ACh in spermatozoa does not have a storage pool [6]. Therefore, synthesis, release, breakdown, and receptor stimulation are closely linked [8]. Dwivedi and Long investigated that the direct and indirect agonist cholinergic agents stimulated the spermatozoa motility, and antagonist cholinergic agents inhibited spermatozoa motility.

Because animal spermatozoa access all cholinergic system compartments, we hypothesized that direct adding ACh increases sperm motility in roosters. Therefore, this study aimed to evaluate the in-vitro effect of the cholinergic agent (ACh) on semen quality in roosters.

## material and methods

### Animals selection and samples collection

This study used ten same-aged indigenous roosters for the sample collection and roosters habituated by abdominal massage for semen collection over one month before starting the experiment. Animals eat a balanced diet (metabolizable energy 2800kcal, crude protein 12%, crude fiber 3.5%, available phosphor 0.35%, calcium 0.7%, salt 1%, and vitamin premix 0.25%, and mineral premix 0.25%) of 90-100g per day. Water was continuously available for it in 16:8 light/dark photoperiods, and the temperature was 20-25°C.

The semen collection takes place twice a week. The semen is collected in a degraded microtube after evaluating the primary semen parameter and pooled together to discards differences between the roosters. Semen with total motility lower than 85% and progressive motility lower than 65% was discarded.

### Extender preparation

Used a Lake extender for rooster semen dilution and shown their components in Table 1. Collected semen was diluted by Lake extender 1:10 ratio and preserved in the refrigerator at 4°C. This is tha Table 1 legend

### Motility evaluation

Sperm motility was Evaluated by light microscopes through a good technician. At slightest five areas, counting a least of 200 sperm were evaluated by technician and assessed Sperm total motility (TM%).

### Viability evaluation

For evaluating spermatozoa viability, was used the eosin-nigrosine stain method. Used the buffer solution to prepare the eosin-nigrosine stain, containing 17.35g sodium glutamate, 1.28g potassium citrate, 8.51g sodium citrate, and 0.68g magnesium chloride in 1L distilled. Dissolve 5g nigrosine with 1g eosin in 100mL of above buffer for preparation eosin-nigrosine stain.

Putting 10uL of eosin-nigrosine stain on the clean slide and added 5uL of diluted semen for it and mixed perfectly by the same pipette. After that, was used another clean slide at 45 degrees to prepare the smear and let it at room temperature to dry. 200-250 spermatozoa under the light microscope at 1000X magnification was counting. The white-colored spermatozoa are life, and all parts of it change their color to pink color have died spermatozoa (Fig 1), finally, expressed the data as a percentage.

**Fig 1.**
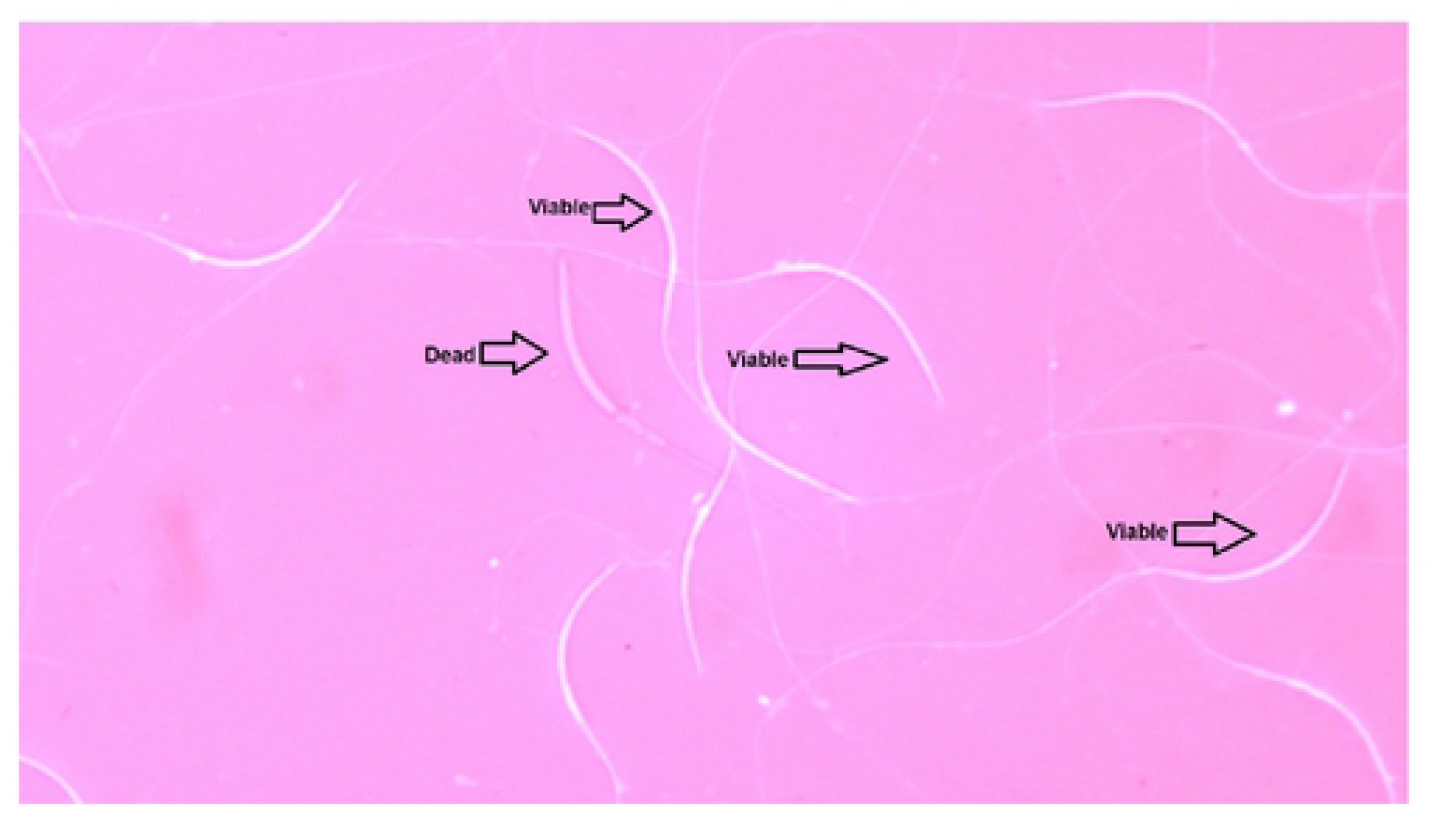
Show the viable and dead rooster spermatozoa by using eosin-nigrosine method at 1000X magnification.

### Acrosome Integrity Evaluation

Evaluated acrosome integrity by using the 4% formaldehyde citrate solution. Added 2.9g tri-sodium citrate dehydrate to 100mL of DW to prepare 2.9% sodium citrate, then 4mL of 2.9% sodium citrate added to 96mL formaldehyde 37% to prepare 4% formaldehyde citrate.

Mixed 10uL of eosin-nigrosine stain, 100uL of 4% formaldehyde citrate, and 5uL diluted semen in 0.5cc microtube, after 30 seconds 15uL of it added to the clean slide and by another clean slide at 45 degrees prepare a smear, and let it for drying at room temperature. Under a light microscope 1000X, magnification enumerates spermatozoa in 10 random fields. Classify tap tip acrosome from others and present it as a percentage; those sperms have tap tip acrosome safe acrosome and vice versa (Fig 2).

**Fig 2.**
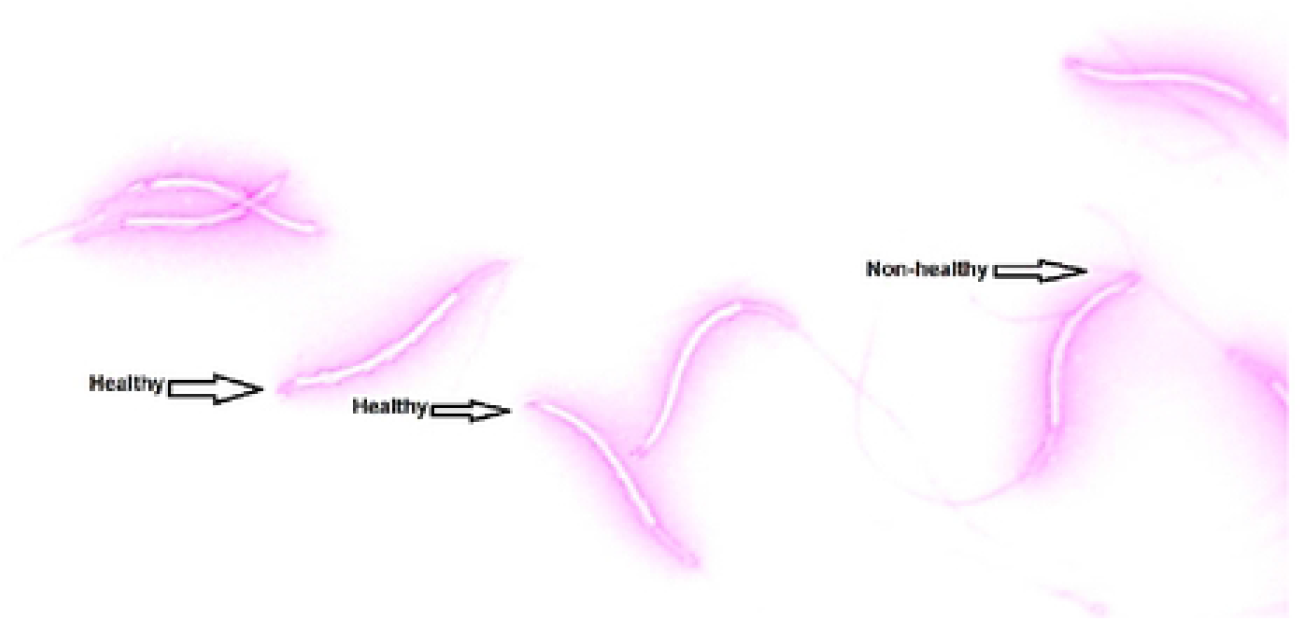
Show the Acrosome integrity by using formalin citrate and eosin-nigrosine stain method at 1000X magnification.

### Plasma membrane integrity evaluation

Used the hypo-osmotic solution to evaluate the plasma membrane integrity. Preparation of the hypo-osmotic solution needs 9 g fructose with 4.9 g sodium citrate dissolved in 1L of distilling water. 10uL of diluted semen added for 100uL of hypo-osmotic solution in 0.5cc micro-tube mixed it and incubated in the water bath for 60 min at 37°C. Added 10uL of incubated sample on a clean slide and mixed with 10uL of eosin-nigrosine stain, finally prepare a smear and late it for drying at room temperature. Seen ten fields under the light microscope at 1000X magnification, non-crawled flagellum spermatozoa are lost their plasma membrane integrity, and vice versa expressed the data as a percentage (Fig 3).

**Fig. 3.**
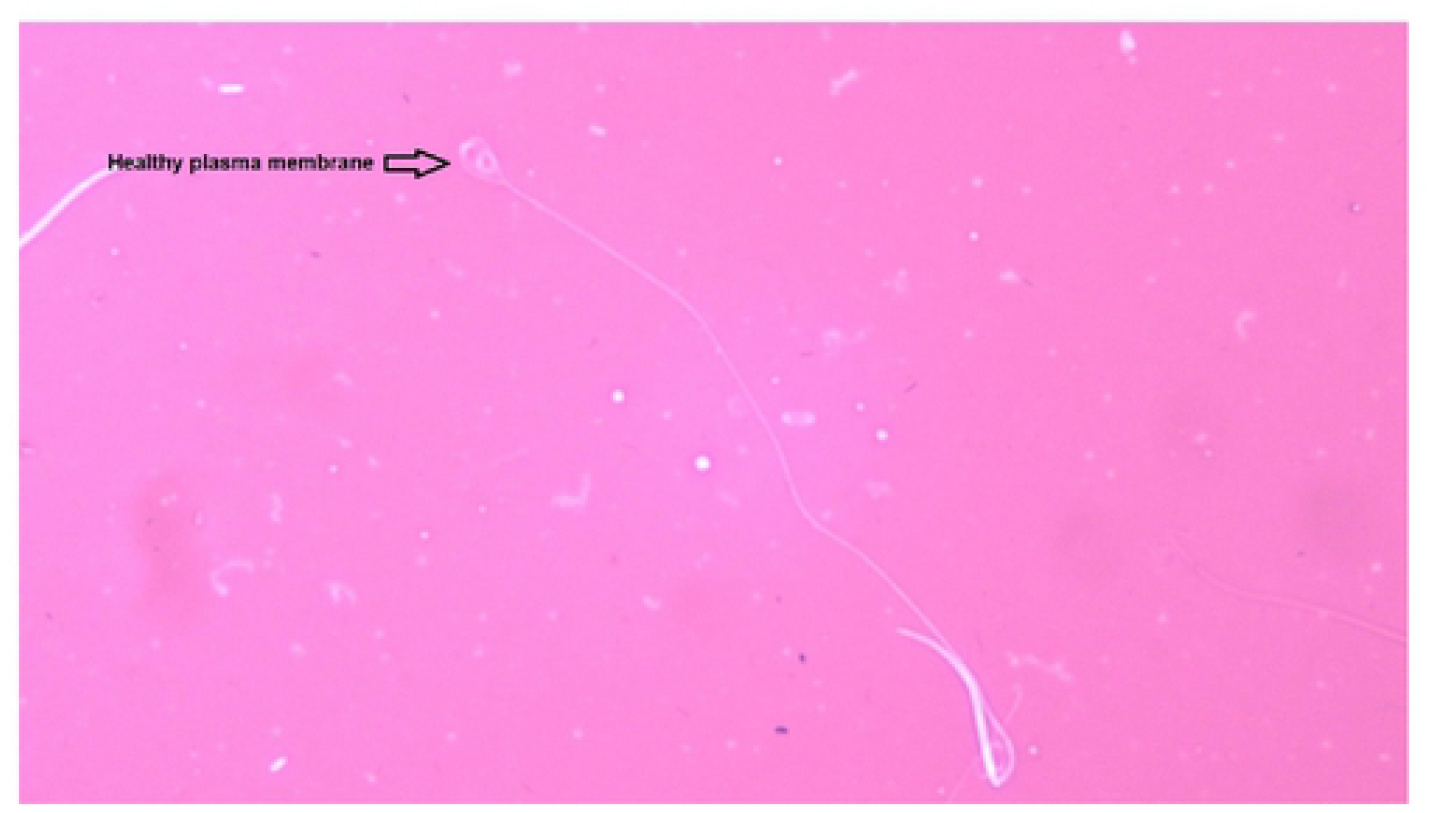
Show the plasma membrane integrity in rooster spermatozoa by using the hypo-osmotic solution test (HOST) technique and using the eosin-nigrosine stain method at 1000X magnification.

### Evaluation of optimal atropine concentration for semen preservation

Tested the acetylcholine in three different concentrations (100μM, 1mM, and 10mM) to determine its optimal and effective concentration for rooster semen recovery. The fresh semen after collection and evaluation motility test, diluted 1:10 ratio by the Lake extender and preserved for 24 hours in at 4°C. Diluted semen by Lake diluent stored for 24 hours at 4°C. Their motility decreased to 50%, added different ACh concentrations, and evaluated their motility at times 0, 5min, 10min 15min, 30min, and 120min at room temperature. Evaluate the viability, acrosome, and plasma membrane integrity for incubated semen with ACh at room temperature after 120min.

### Statistical analysis

Used the utterly random design with ten replicate, and the obtained data were submitted to analysis of variance using the SAS statistical package’s general linear model (GLM) procedure (SAS Institute, 2002). Duncan’s multiple range test procedures detected significant differences among means of treatments. The differences were considered significant at P < 0.05.

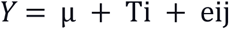

Where Y is the measured parameter, μ is the mean of the parameter, T_i_ is the effect of the treatment, and e_ij_ is the effect of the errors.

## Result and discussion

The movements of spermatozoa appear to be sensitive to various agents that affect synaptic transmission [9]. Flagella are responsible for spermatozoa movements and contractile proteins [4]. Contractile proteins of flagella are generally similar to the smooth muscle. Scientists believe that the ACh cycle is responsible for the contraction and relaxation of flagella [5]. In the animal spermatozoa, all compartments of the cholinergic system include ACh, ChAT enzyme [6], AChE [7], and both nAChRs and mAChRs are present [5]. Binding the ACh to their receptors activates them and causes membrane depolarization through the influx of Na^+1^ and Ca^+2^ [10]; after the membrane depolarization, it increases sperm motility by increasing flagella beating [11]. Therefore, as shown in table 1, after adding ACh at 10mM concentration for nearly 50%, motile rooster sperms significantly increase motility to almost 80%. Sperm motility after adding ACh for it their motility recovered to nearly 80%. It remained in this range of motility to 30min and after that decreased it (Fig 4). After two hours of recovery, sperm viability compared to control reduced significantly (Fig 5). This decrease of viability belongs to the high motility of spermatozoa during these two hours compared to the motility of the control group [12], but adding ACh does not affect the sperm acrosome and plasma membrane integrity (Figs 6 and 7).

**Fig 4.**
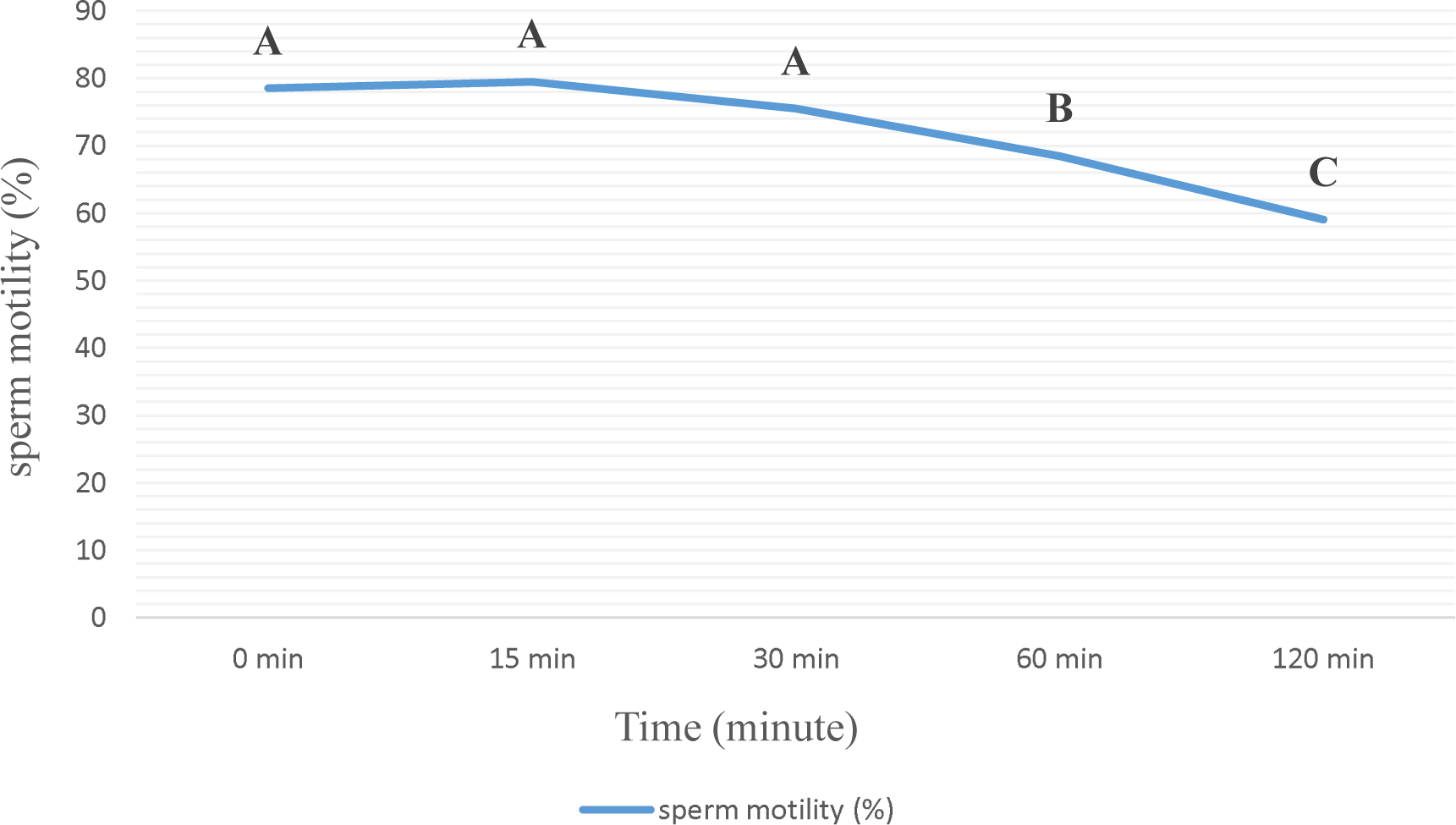
Total sperm motility during 2 hours of incubation at room temperature after recovery by adding 10mM ACh for 24 hours preserved semen at 4°C.

**Fig 5.**
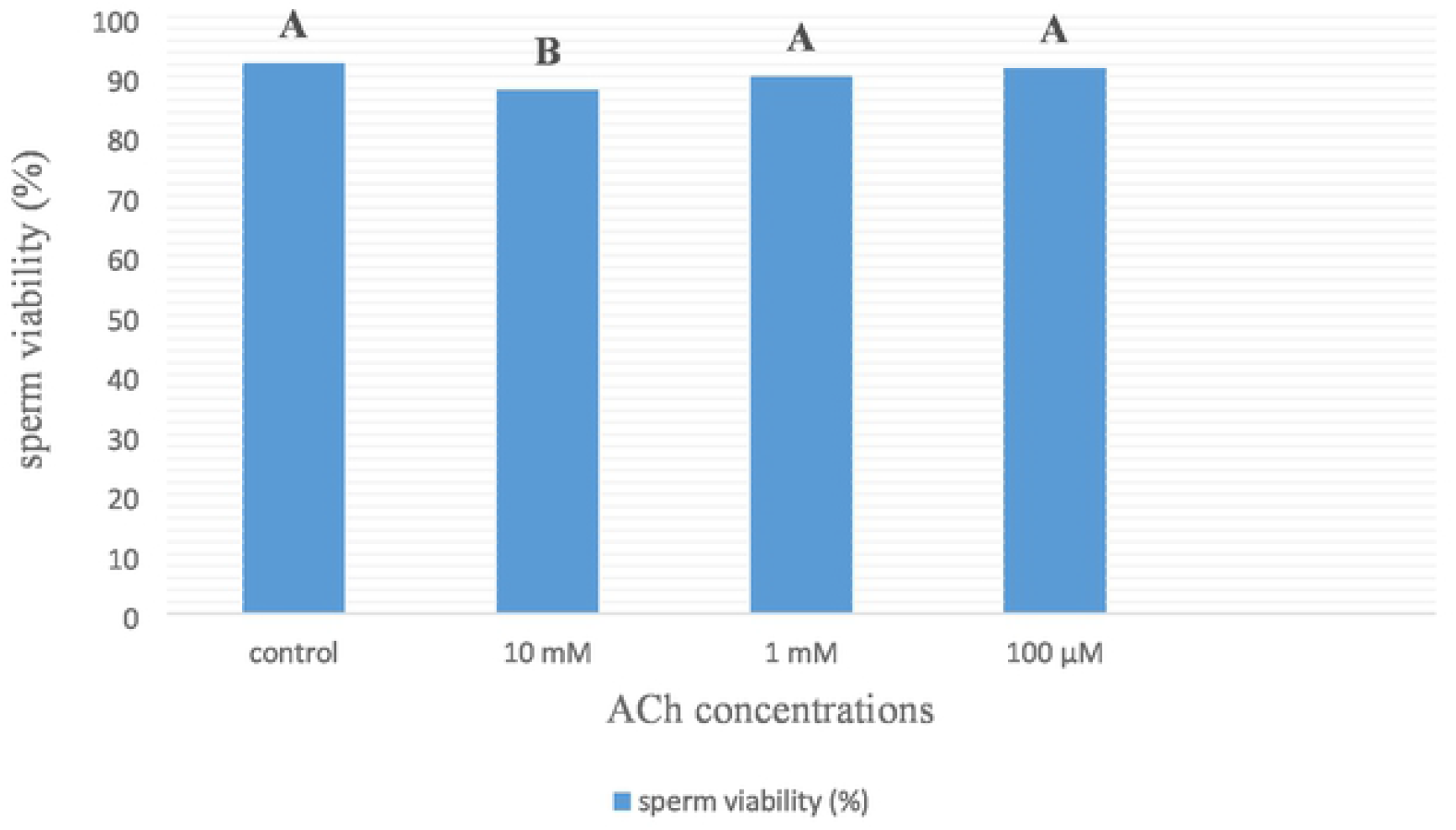
Sperm viability (%) after 2 hours’ storage at room temperature from their recovery by adding different ACh concentration for 24 hours preserved semen at 4°C.

**Fig 6.**
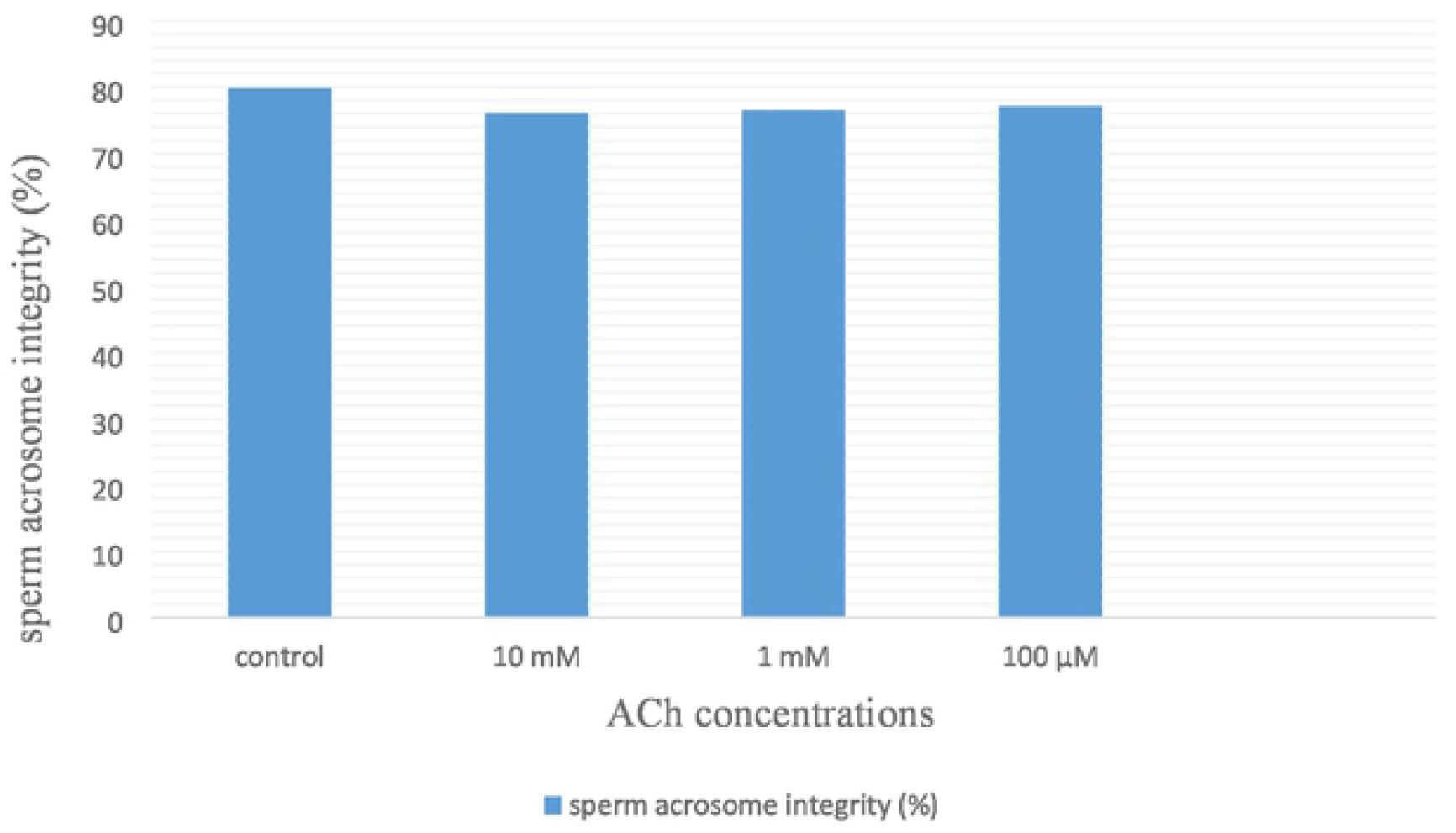
Sperm acrosome integrity (%) after 2 hours’ storage at room temperature from their recovery by adding different ACh concentration for 24 hours preserved semen at 4°C.

**Fig 7.**
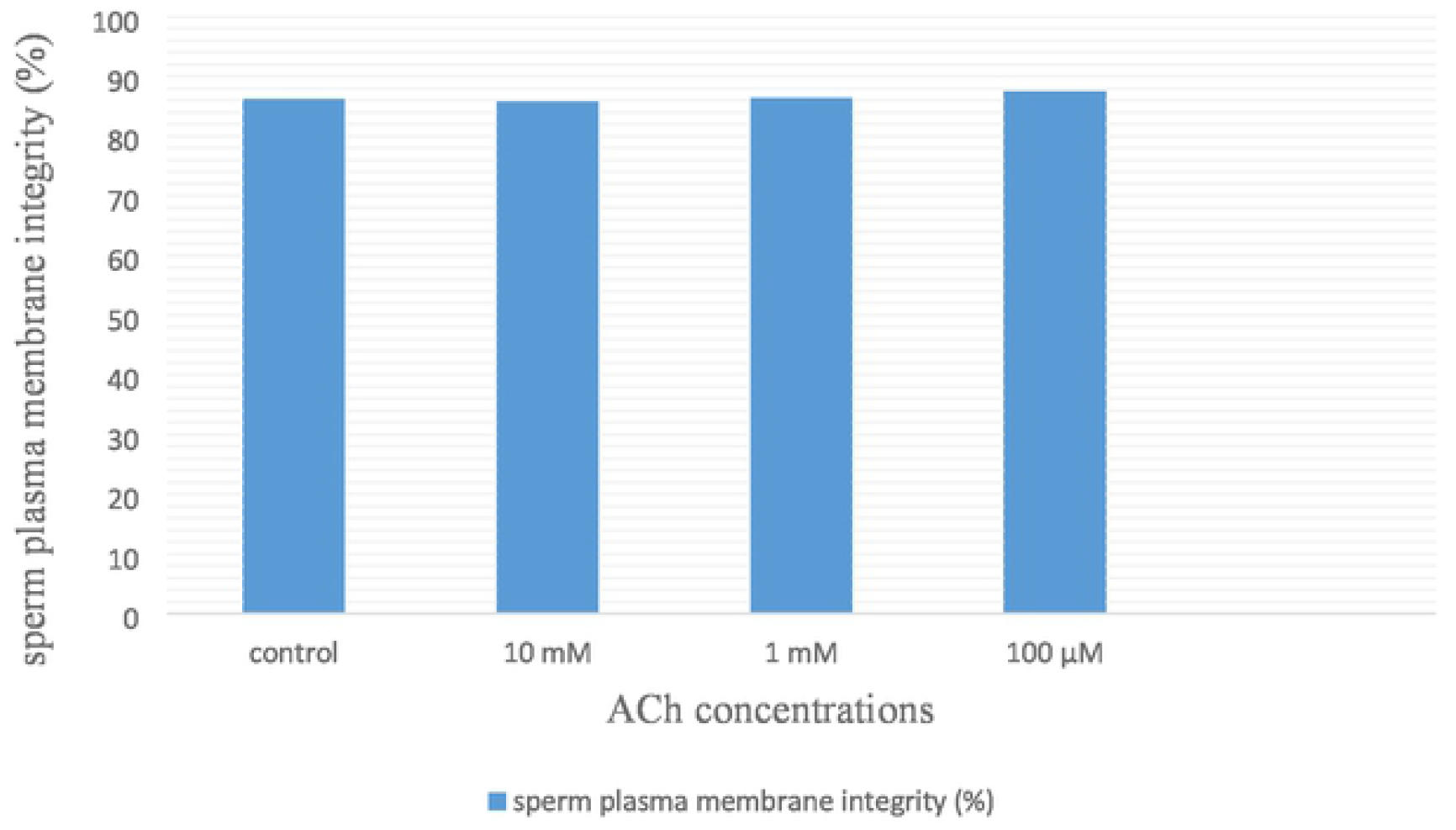
Sperm plasma membrane integrity (%) after 2 hours’ storage at room temperature from their recovery by adding different ACh concentration for 24 hours preserved semen at 4°C.

By adding ACh to attenuate spermatozoa, it recovered to an active form. This recovery belongs to stimulating cholinergic receptors (muscarinic and nicotinic) on the spermatozoa plasma membrane at all parts but is more distributed at the neck and midpiece region near the flagella, therefore activation of the muscarinic and nicotinic receptor produces an action potential in the plasma membrane and activate several intracellular pathways to increase intracellular calcium concentration [13–17) see figs 8 and 9. Therefore, an increase in intracellular calcium leads to increased production of ATP by mitochondria through altering the sensitivity of calcium-sensitive mitochondrial matrix enzymes [18]. Likewise, an increase in the intracellular calcium concentration leads to activation of retro flow movement in the intraflagellar transport. This activation of intraflagellar flow leads to bent the flagella and flagella beating [19]. Therefore, adding ACh can increase sperm movement by increasing ATP production through mitochondria (using it for cellular metabolism) and increasing flagellar beating. Also, according to Figs 5-7, ACh from 100μM to 10mM concentration does not have any cytotoxicity for rooster spermatozoa and can be used as a safe treatment.

**Fig 8.**
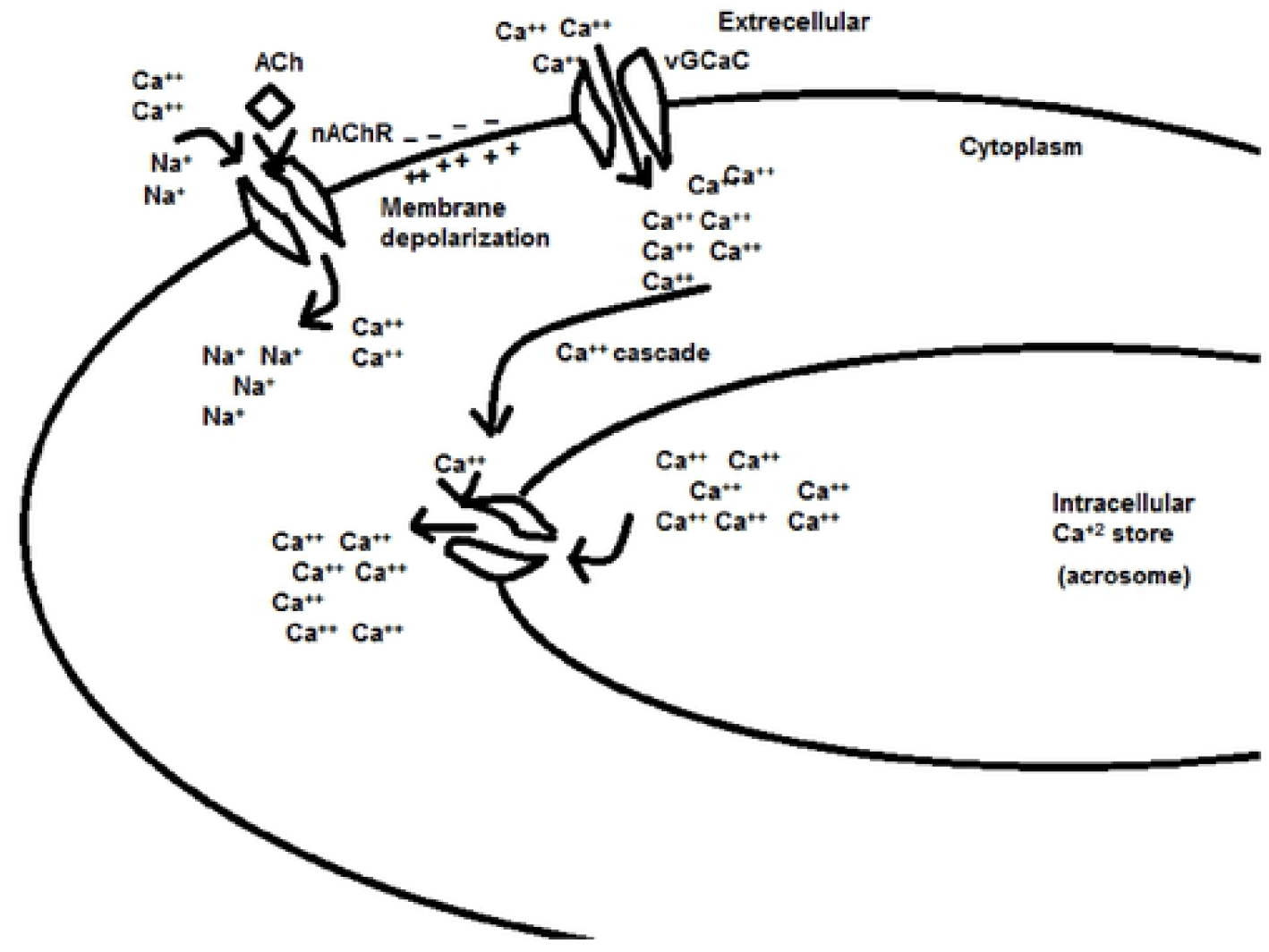
Mechanism action of ACh through the nicotinic cholinergic receptors.

**Fig 9.**
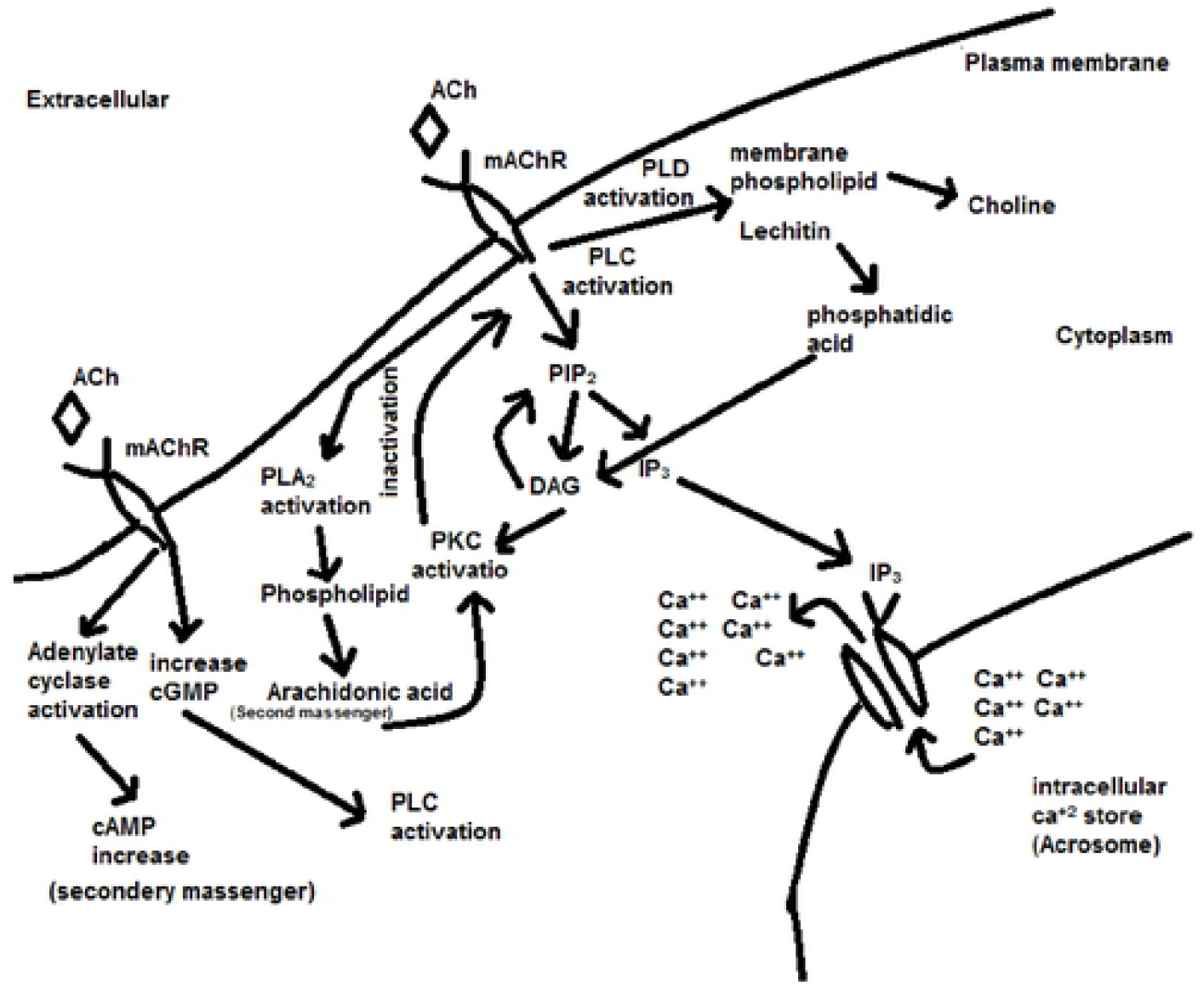
Mechanism action of ACh through the muscarinic cholinergic receptors.

## conclusion

The present study showed that adding acetylcholine to the rooster semen at 100μM to 10mM concentration does not have any side effect or toxicity for rooster semen and is entirely safe. Also, adding 10mM acetylcholine to low motile rooster semen (50%) can increase and recover motility of the spermatozoa and remained this increase of motility (nearly 75%) for 30min more. Therefore, this finding can help the parent stock breeders to use the low motile semen from their roosters when it decreases the semen quality and is used in the artificial insemination technique.

## Supporting information

Table 1: Effect of different acetylcholine concentrations on sperm motility during two hours (h) at room temperature after 24 hours (h) storage at 4oC

