## Supplementary material for "An in-vitro investigation of effects of acetylcholine on attenuate and low motile diluted rooster semen quality": Table 1: Effect of different acetylcholine concentrations on sperm motility during two hours (h) at room temperature after 24 hours (h) storage at 4oC

| Table 1: **Effect of different acetylcholine concentrations on sperm motility during two hours (h) at room temperature after 24 hours (h) storage at 4^o^C.** | | | | |
| --- | --- | --- | --- | --- |
| **Time** | **Control** | **10mM** | **1mM** | **100μM** |
| **Sperm motility** | | | | |
| **0 min** | 50 ± 4^b^ | 78.5 ± 1^a^ | 58.5 ± 4^b^ | 52.5 ± 5^b^ |
| **15 min** | 28 ± 3^bc^ | 79.5 ± 1^a^ | 32.5 ± 4^b^ | 21 ± 3^c^ |
| **30 min** | 20 ± 4^b^ | 75.5 ± 2^a^ | 20 ± 3^b^ | 9 ± 2^c^ |
| **1 h** | 12.5 ± 2^b^ | 68.5 ± 1^a^ | 17 ± 3^b^ | 5.5 ± 3^b^ |
| **2 h** | 2.5 ± 1^b^ | 59.0 ± 2^a^ | 4 ± 1^b^ | 1 ± 1^b^ |
| Different superscripts within the same row indicate significant differences among groups (P<0.05).  Lake extender without acetylcholine (control), Lake extender with 10mM acetylcholine (10mM), Lake extender with 1mM acetylcholine (1mM), Lake extender with 100μM acetylcholine (100μM). | | | | |
